# Identifying cell type specific driver genes in autism-associated copy number loci from cerebral organoids

**DOI:** 10.1101/2020.11.15.375386

**Authors:** Elaine T. Lim, Yingleong Chan, Mannix J. Burns, Xiaoge Guo, Serkan Erdin, Derek J.C. Tai, Julia M. Reichert, Ying Kai Chan, Jessica J. Chiang, Katharina Meyers, Xiaochang Zhang, Christopher A. Walsh, Bruce A. Yankner, Soumya Raychaudhuri, Joel N. Hirschhorn, James F. Gusella, Michael E. Talkowski, George M. Church

## Abstract

Neuropsychiatric and neurodevelopmental disorders have been particularly challenging to study using animal models, and recently, human-derived cerebral organoids demonstrate great promise for discovering molecular processes that are important in these disorders. However, several challenges remain in achieving robust phenotyping to discover cell type specific genes. We perform RNA sequencing on 71 samples comprising of 1,420 cerebral organoids from 25 donors, and describe a framework (Orgo-Seq) to identify cell type specific driver genes, for 16p11.2 deletions and 15q11-13 duplications. We identify neuroepithelial cells as critical cell types for 16p11.2 deletions, and discover novel and previously reported cell type specific driver genes. Finally, we validated our results that mutations in *KCTD13* in the 16p11.2 locus lead to imbalances in the proportion of neuroepithelial cells, using CRISPR/Cas9-edited mosaic organoids. Our work presents a quantitative discovery and validation framework for identifying cell type specific driver genes associated with complex diseases using cerebral organoids.

## MAIN

Recent advances in three-dimensional brain models differentiated from human induced pluripotent stem cells (iPSCs), also known as cerebral organoids, have shown that these *in-vitro* systems can mimic molecular and cellular processes in the developing human fetal brain^1,2^. Cerebral organoids also show great promise for modeling processes that are perturbed in neurodevelopmental and neuropsychiatric disorders such as microcephaly and autism spectrum disorders (ASD)^2,3^. These human-derived organoids have significant advantages over animal models, and in many instances, might prove to be complementary to animal models for studying complex human neurological diseases.

For instance, copy number variants (CNVs) in 16p11.2 and 15q11-13 have been robustly associated with ASD^4^, but it is unclear what cell types are affected by these CNVs, and which of the genes in these CNV loci are driver genes leading to ASD-associated phenotypes when mutated. Clinical studies have shown that individuals with 16p11.2 deletions have increased brain sizes, while individuals with 16p11.2 duplications have decreased brain sizes^5,6^. A systematic dissection of all genes in the 16p11.2 locus using brain sizes as the phenotypic readout in zebrafish identified *KCTD13* as a key driver gene modulating the proportion of neural progenitor cells^7^. However, recent studies in mice and zebrafish with deleted *KCTD13* did not observe increased brain sizes or neurogenesis in these mutant mice and zebrafish^8,9^.

Given the vast differences between the human brain and brains from animal models, human post-mortem brain tissue could ideally be used to resolve such conflicting results from animal models. However, ASD is a genetically heterogeneous complex disorder and post-mortem brain samples from individuals with specifically 16p11.2 deletions or duplications are not easily available currently^10^. To address these questions, we sought to identify the critical cell type(s) and cell type specific driver gene(s) in the 16p11.2 locus using patient-derived human cerebral organoids. We further apply our approaches to patient-derived cerebral organoids with 15q11-13 duplications where there is less definitive evidence for critical cell types and candidate driver genes for the 15q11-13 duplication locus.

One key challenge remains in using cerebral organoids as a model system for understanding complex neurological disorders. Prior literature has demonstrated that the cerebral organoids comprise of many different cell types found in the human brain, and individual organoids can be heterogeneous in their cell type compositions detected using single-cell RNA sequencing^1^. This poses additional challenges for detecting robust cellular and molecular differences between cerebral organoids differentiated from individuals with different genetic backgrounds. To address this key challenge, we differentiated a large number of 1,420 organoids from 25 individuals with diverse backgrounds (71 samples with 20 organoids per sample), to systematically quantify and identify the inherent variability in whole-transcriptome bulk RNA sequence data derived from the organoids.

A second key challenge is to robustly detect cell type specific driver genes that are perturbed in organoids differentiated from patients compared control individuals. One advantage of using single-cell RNA sequencing is that we can perform unbiased discovery of critical cell types associated with diseases. However, single-cell RNA sequencing technologies capture only 10-20% of all transcripts^11^, and the expression of many disease-associated genes, such as most of the genes in the 16p11.2 locus, might not be detectable with single-cell RNA sequencing^1^.

Here we developed a novel, quantitative phenotyping framework (termed Orgo-Seq for “<u>**Org**</u>an<u>**o**</u>id <u>**Seq**</u>uencing”, **Fig. 1**), which allows researchers to identify cell type specific driver genes associated with complex diseases, by using bulk RNA sequence data from patient-derived organoids, and at the same time, utilizing single-cell RNA sequence data as an input reference panel. This allows us to overcome the capture limitation with single-cell RNA sequencing to discover cell type specific driver genes. We applied Orgo-Seq for two ASD-associated CNVs in the 16p11.2 and 15q11-13 loci, by using previously published single-cell RNA sequence data that has been generated from cerebral organoids^1^ and existing bulk RNA sequence data from the BrainSpan Project^12^. Finally, we described a mosaic cerebral organoid framework using CRISPR/Cas9 editing to validate one of our key findings from Orgo-Seq for the 16p11.2 locus. Our work presents a quantitative framework to identify cell type specific driver genes in a complex disease using bulk RNA sequencing on patient-derived cerebral organoids, and a CRISPR/Cas9 based mosaic cerebral organoid system to validate the findings from the patient-derived cerebral organoids.

**Figure 1:**
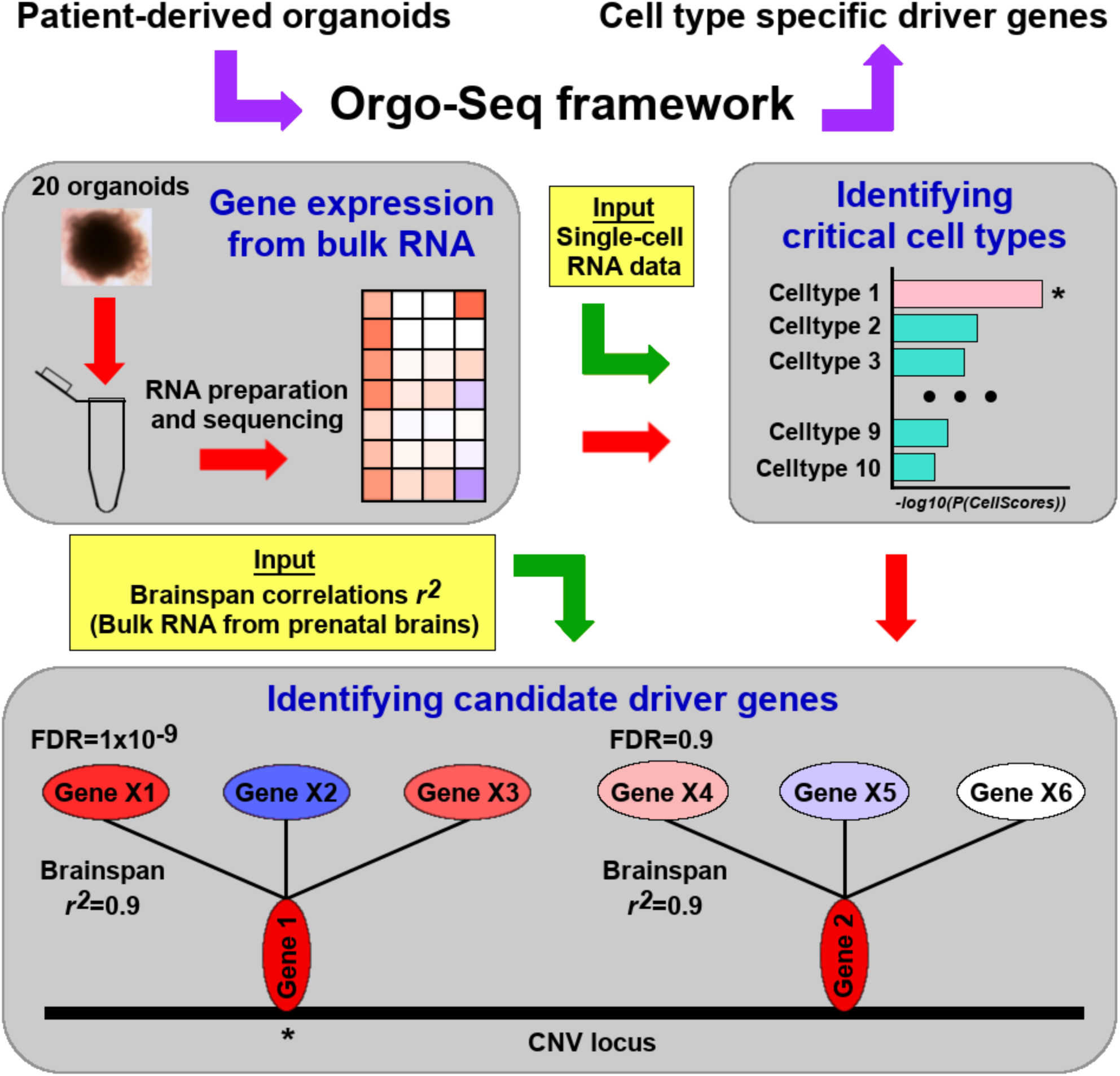
Orgo-Seq framework to identify critical cell types and driver genes. Figure illustrating the Orgo-Seq framework, which is a bridge between patient-derived organoids and understanding disease biology through the discovery of cell type specific driver genes.

## RESULTS

### Identifying variability in RNA sequence data from cerebral organoids

To evaluate the extent of heterogeneity in cerebral organoids, it is important to first identify the sources of variability in RNA sequence data from organoids differentiated from a large number of donors. We obtained iPSCs from 12 control donors (termed “controls”) and 13 donors with 16p11.2 deletions or 15q11-13 duplications (termed “cases”). DNA was extracted from the iPSCs (**Supplementary Table 1**) and CNV detection was performed on iPSCs from all donors using array comparative genomic hybridization or aCGH (**Supplementary Table 2**) and whole-exome sequencing was performed to detect smaller exonic CNVs (**Supplementary Table 3**). All controls were confirmed not to harbor any CNVs within the two ASD-associated loci in 16p11.2 and 15q11-13.

We differentiated cerebral organoids using the 25 iPSCs from the cases and controls for 46 days, by adapting a previously described method^13^ (**Supplementary Fig. 1**). For each individual, we performed RNA sequencing on 1 to 3 replicates of their cerebral organoids, resulting in a total of 71 samples (**Supplementary Table 1**). We compared the standard deviations in gene expression between replicates for each individual (intra-individual), as well as across organoids differentiated from different individuals (inter-individual), and found that there were 860 genes (7.6% of all expressed genes) that showed high intra-individual variability, and 869 genes (7.7% of all expressed genes) that showed high inter-individual variability (**Supplementary Fig. 2A-B**). These genes with high intra-individual or inter-individual variability were enriched in processes involved in nervous system development, neurogenesis and cell differentiation (**Supplementary Table 4**), which might contribute to the inherent variability in spontaneous differentiation of these cerebral organoids. While these genes might be important in neuropsychiatric disorders such as ASD, the technical variability in expression for these genes is high. For our downstream analyses, we removed these highly variable genes and focused on a smaller, robust group of genes with low technical variability in expression, and there are 9,978 such unique genes that were detected in the organoids (**Supplementary Tables 5-8**).

To quantify the variability in bulk RNA sequence data from the cerebral organoids, we calculated pairwise Pearson’s correlations (*r*^2^), and the histograms of the Pearson’s correlations are shown in **Supplementary Figs. 2C-H**. When we calculated the pairwise Pearson’s correlations between replicates from the same individual (intra-individual), across all control samples after removing the outlier genes, we observed high mean and minimum *r*^2^ of 0.97 and 0.89 respectively (**Supplementary Fig. 2C**). The mean and minimum intra-individual *r*^2^ when comparing all samples (cases and controls) were similarly high at 0.97 and 0.88 respectively (**Supplementary Fig. 2D**). Prior to removing the outlier genes, the mean and minimum intra-individual *r*^2^ across all samples were slightly lower at 0.96 and 0.84 respectively (**Supplementary Fig. 2E**).

When we calculated all pairwise Pearson’s correlations between samples from different control individuals (inter-individual) after removing the outlier genes, we observed high mean and minimum *r*^2^ of 0.94 and 0.71 respectively (**Supplementary Fig. 2F**). The mean and minimum inter-individual *r*^2^ when comparing all samples (cases and controls) were similar at 0.94 and 0.71 respectively (**Supplementary Fig. 2G**). Prior to removing the outlier genes, the mean and minimum inter-individual *r*^2^ across all samples were slightly lower at 0.92 and 0.69 respectively prior to removing the outlier genes (**Supplementary Fig. 2H**).

All the intra-individual and inter-individual correlations were high, demonstrating that there is low inherent variability in bulk RNA sequence data from the cerebral organoids using our approach. Similar to a previous report, we observed higher intra-individual correlations than inter-individual correlations, confirming that whole transcriptome data from the cerebral organoids can reflect biological differences between individuals that are not due to technical differences between replicates differentiated from the same individual^3^.

Principal components analyses on all case and control samples showed that most of the variance in gene expression (88%) can be accounted for by the first principal component (PC1) alone (**Supplementary Fig. 3A**). We further explored several factors that might influence variability in gene expression from the cerebral organoids, such as ethnicity, sex, age, origin of sample used for iPSC reprogramming and type of reprogramming (**Supplementary Fig. 3B-G**). We performed correlation tests between the first 3 principal components and observed that age, origin of sample and the type of reprogramming are significantly correlated with PC1 alone, but not with the second and third principal components (**Supplementary Fig. 4**). These results suggest that PC1 is a surrogate variable for age, origin of sample and type of reprogramming, and subsequently, we included PC1 as a covariate in the differential expression analyses.

### RNA sequence data from cerebral organoids show highest correlation with prenatal post-mortem brain samples

To determine how accurately RNA sequence data from cerebral organoids can reflect RNA sequence data from post-mortem human brain samples, we compared our data with data from 578 post-mortem brain samples in the BrainSpan Project^12^. We observed a wide range of correlations (Pearson’s *r*^2^ from 0.21 to 0.82, **Supplementary Fig. 5A**). The mean correlations between the organoids with individual prenatal brain samples (maximum *r*^2^=0.67, minimum *r*^2^=0.38) were significantly higher than the mean correlations between the organoids with individual postnatal brain samples (maximum *r*^2^=0.58, minimum *r*^2^=0.28), shown in **Supplementary Fig. 5B** (two-sided Wilcoxon P<2.2×10^−16^).

The highest mean correlations were with prenatal brain samples from regions such as the hippocampus (*r*^2^=0.61), primary visual cortex (*r*^2^=0.61) and amygdala (*r*^2^=0.61), shown in **Supplementary Fig. 5C**. The lowest correlations with prenatal brain samples were across brain regions such as the mediodorsal thalamus (*r*^2^=0.49), ventrolateral prefrontal cortex (*r*^2^=0.56) and primary somatosensory (*r*^2^=0.56). Similar observations were made when comparing RNA sequence data from cerebral organoids prior to removing the outlier genes (**Supplementary Fig. 5D-F**). These observations suggest that RNA sequence data from the cerebral organoids are highly correlated with RNA sequence data from prenatal brain samples but not with postnatal brain samples.

### Transcriptome data in cerebral organoids accurately reflect copy number changes

Given that RNA sequence data from the cerebral organoids are highly correlated with RNA sequence data from prenatal post-mortem brain samples, and in the absence of fetal post-mortem brain samples with specific mutations such as CNVs in 16p11.2, we can effectively use cerebral organoids as a model system for studying mutation-specific transcriptomic processes that are important in human neurodevelopmental diseases. The 16p11.2 locus encompasses 22 genes that are expressed in the organoids. In our study, there are 3 individuals with ASD and 16p11.2 deletions (whom we termed as “probands”), 6 individuals with 16p11.2 deletions but are not clinically diagnosed with ASD (whom we termed as “resilient” individuals), and 12 control unaffected individuals without 16p11.2 deletions. For differential expression analyses, we used linear regression with the first principal component as a covariate, and performed multiple hypotheses correction using the Benjamini-Hochberg false discovery rate (FDR). We further checked the first 2 principal components, but did not observe major stratification between the cases and controls (**Supplementary Fig. 6**).

We performed three sets of differential expression analyses on RNA sequence data from cerebral organoids differentiated from these individuals. **SetA** comparing all 9 individuals with 16p11.2 deletions with 12 control individuals without 16p11.2 deletions (**Supplementary Table 5**); **SetP** comparing the 3 probands with ASD and 16p11.2 deletions with 12 control individuals without 16p11.2 deletions (**Supplementary Table 6**); and **SetD** analyses comparing only the individuals with 16p11.2 deletions: 3 probands with 16p11.2 deletions versus 6 resilient individuals with 16p11.2 deletions (**Supplementary Table 7**). We observed 2,681 genes with FDR≤0.05 in the SetA comparison, and 1,853 genes with FDR≤0.05 in the SetP comparison.

If RNA sequence data from the cerebral organoids can accurately reflect the underlying genetic mutations in the DNA (hemizygous deletions in the 16p11.2 locus or duplications in the 15q11-13 locus), then we should expect to observe that many of the genes in the 16p11.2 locus are down-regulated with fold changes of ∼0.5 in the cases compared to controls (**Supplementary Tables 5-6**). For the SetA comparison, 19 of the 22 genes in the 16p11.2 locus (excluding *SULT1A4* [MIM 615819], *SULT1A3* [MIM: 600641] and *QPRT* [MIM: 606248]) are significantly differentially expressed with FDR≤0.05 (**Supplementary Table 5**). The average fold-change for the 19 significantly differentially expressed genes in the 16p11.2 locus in the SetA comparison is 0.73, which closely reflects the 0.5-fold change in copy number across the locus. 17 of these 19 genes are also significantly differentially expressed in the smaller SetP comparison, with an average fold-change of 0.64 (**Supplementary Table 6**).

Many risk loci associated with complex disorders such as ASD have incomplete penetrance^14^, and are present in control individuals who are not clinically diagnosed with these disorders, albeit at a lower prevalence than in affected individuals. A two-hit model was previously described for another microdeletion locus in 16p12.1, where the microdeletion was reported to exacerbate neurodevelopmental phenotypes in association with other large CNVs^15^. We asked if we could similarly detect a second genetic factor that contributes to increased risk for ASD, in addition to the 16p11.2 deletion background, and this might explain for the incomplete penetrance observed with the 16p11.2 deletions. The SetD comparison between individuals with ASD and 16p11.2 deletions versus resilient individuals with 16p11.2 deletions did not yield any gene with FDR≤0.05. There is less than 80% power to detect a gene with small effects (fold changes of less than 4) at FDR≤0.05 given our current sample sizes (**Supplementary Table 8**). Larger numbers of individuals with 16p11.2 deletions will be needed to identify a second genetic hit with small effects, or it might be possible that the second hit is driven by non-genetic factors or by genetic factors that are not expressed in the cerebral organoids. However, we can exclude the hypothesis that there is a second genetic hit with large effects given our current sample sizes.

Out of the 25 individuals in our study, there are 4 individuals with ASD and 15q11-13 duplications, and 12 control unaffected individuals without 15q11-13 duplications, and we similarly performed whole-transcriptome RNA sequencing on cerebral organoids differentiated from these individuals in triplicates (**Supplementary Table 1**). There are 16 genes that are significantly differentially expressed in the individuals with ASD and 15q11-13 duplications versus unaffected control individuals with FDR ≤ 0.05 (**Supplementary Table 9**). Out of the 16 genes, 5 of them are found in the 15q11-13 locus (*HERC2* [MIM 605837], *TUBGCP5* [MIM 608147], *CYFIP1* [MIM 606322], *NIPA2* [MIM 608146] and *UBE3A* [MIM 601623]). The average fold-change for the 5 genes in the 15q11-13 locus that are significantly differentially expressed is 1.48, which closely reflects the 1.5-fold change in copy number across the locus, suggesting that the RNA sequence measurements are robust and quantitative. Three other genes in the 15q11-13 locus (*OCA2* [MIM 611409], *NIPA1* [MIM 608145] and *GABRB3* [MIM 137192]) had fold-changes of greater than 1, but were not significantly differentially expressed in the organoids from the individuals with ASD compared to unaffected controls. Another 5 genes in the 15q11-13 locus (*SNURF* [MIM 182279], *SNRPN* [MIM 182279], *NDN* [MIM 602117], *IPW* [MIM 601491] and *MAGEL2* [MIM 605283]) had fold-changes of ∼1, possibly because of epigenetic imprinting at the locus. We did not detect smaller duplications that might encompass only a subset of these genes in the 15q11-13 locus for these individuals with ASD, using aCGH and whole-exome sequencing (**Supplementary Tables 2-3**).

### Deconvolution of bulk RNA sequence data identifies critical cell types for 16p11.**2 deletions and 15q11-13 duplications**

Identifying the cell types that are perturbed as a result of mutations in disease-associated loci allows us to perform direct experiments on the relevant cell types to understand molecular processes that are important in disease. Moreover, identifying perturbations in these critical cell types can highlight cellular endophenotypes for screening therapeutic targets using cerebral organoids. A recent publication had identified 10 major clusters of cell types (c1-10) in cerebral organoids using single-cell RNA sequencing (Drop-seq), as well as a list of genes that tag each of the 10 clusters^1^. We separated the lists of genes into a set of cell type specific genes (ranging from 47 to 266 genes; **Supplementary Table 10**) that uniquely identifies each cluster of cell types, and a set of non-cell type specific genes that are found in multiple clusters (ranging from 12 to 49 genes; **Supplementary Table 10**).

Using these genes as a reference panel for defining cell types in cerebral organoids, we evaluated if the differentially expressed genes identified from our bulk RNA sequence data between the cases and controls, are preferentially enriched for cell type specific genes in any of the 10 cell types (**Fig. 1**). We developed a statistic termed *CellScore*, which is the difference between the weighted sum of all cell type specific genes and the weighted sum of all non-cell type specific genes for each cluster of cell types, and the weights are the *-log*_*10*_(P-values) from our differential expression results in cerebral organoids. This allows us to identify transcriptomic signatures arising from the cell type specific genes for each cluster, rather than the non-cell type specific genes contributing to multiple clusters. Next, we evaluated the significance of our observed *CellScores* using permutations (**Supplementary Fig. 7**).

When we applied the *CellScore* evaluation to the differential expression results from the 16p11.2 SetA comparison, we found that the cell cluster comprising of mainly neuroepithelial cells (c9) and unknown cell cluster (c6) are significantly perturbed (P(*CellScore*) = 5.3×10^−3^ and P(*CellScore*) = 1.6×10^−4^ respectively, **Fig. 3A, Supplementary Table 11**). When we applied the *CellScore* evaluation to the differential expression results from the 15q11-13 organoids, we found that there were no cell clusters that were significantly perturbed with a threshold of P(*CellScore*) ≤ 0.01, although the top cell cluster identified was the stem cell cluster (c10), with P(*CellScore*) = 0.017, shown in **Fig. 3B** and **Supplementary Table 11**. Our results suggest that neuroepithelial cells (c9) and the unknown cell type (c6) are critical cell types perturbed in the 16p11.2 organoids, and stem cells (c10) are potential critical cell types perturbed in the 15q11-13 organoids.

### Deconvolution of bulk RNA sequence data on post-mortem brain samples validates top critical cell type identified from cerebral organoids

A prior publication had performed RNA sequencing on post-mortem brain samples of cortex that were obtained from 9 individuals with 15q11-13 duplications and 49 control individuals^10^. We calculated *CellScores* for each of the 10 cell type clusters using the differential expression results from the post-mortem brain samples, and calculated a weighted average P(*CellScore*) using the results from the patient-derived cerebral organoids and post-mortem brain samples with 15q11-13 duplications (**Supplementary Table 12**). Similar to our results from the patient-derived cerebral organoids, there were no cell type clusters identified from the post-mortem brain samples that was significantly perturbed with P(*CellScore*) ≤ 0.01, although the top cell type cluster identified was still the stem cell cluster (c10), with weighted average P(*CellScore*) = 0.03.

### Non-cell type specific co-transcriptional network modeling cannot prioritize driver genes in 16p11.**2 and 15q11-13**

These ASD-associated CNVs are typically large and span across at least 10 genes. Similar to the identification of driver versus passenger genes in cancers, it has been challenging to identify which of the genes in these ASD-associated CNV loci are more likely to be driver genes. The prioritization of candidate driver genes is important for follow-up studies, for instance, to create single-gene knockouts in animal models or organoids for understanding the biological effects of knockouts in these genes.

In the 16p11.2 locus, a prominent study using zebrafish identified *KCTD13* [MIM 608947] as the key causal gene in the locus^7^, although other studies have also shown strong evidence for other genes in the locus such as *TAOK2* [MIM 613199] and *MAPK3* [MIM 601795]^16-18^. CNV analyses on the whole-exome sequence data from one ASD proband with 16p11.2 deletion in our study (14824.x13) found a smaller exonic deletion spanning across exons in *TAOK2* and an intron in *BOLA2B* [MIM 613182] (**Supplementary Table 3**).

In the 15q11-13 locus encompassing 11 genes, several studies have identified *UBE3A* as the major causal gene for ASD^19,20^, although there is supportive evidence for other candidate causal genes such as *CYFIP1* and *HERC2* in the locus^21,22^. Whole-exome sequencing on the iPSCs from one of the ASD probands with 15q11-13 duplication (901) and her unaffected mother (902) showed that they harbored a rare stop-gained mutation (p.Q3441X) in *HERC2*, which is one of the genes in the 15q11-13 locus.

The expression for the genes in the 16p11.2 and 15q11-13 loci range from the 1.8^th^ to 91^st^ percentiles detected from bulk RNA sequencing (**Supplementary Table 13**), and the expression for most of these genes cannot be detected from sequencing a relatively small number of cells using single-cell RNA sequencing^1^. We hypothesized that bulk RNA sequence from the patient-derived cerebral organoids can be harnessed to identify candidate driver genes in these CNV loci. One approach for evaluating the functional effects of genes is to quantify the effects of transcriptomic perturbations in the cerebral organoids. Our assumption is that candidate driver genes are likely to result in more perturbations in downstream genes than the candidate passenger genes. To identify downstream targets of each gene in an unbiased manner, we first calculated the Pearson’s correlations for each of the genes of interest in the CNV loci, with all genes detected from RNA sequencing in the BrainSpan Project, and used the correlations in expression from the BrainSpan Project as a proxy for co-expression connectivity with our genes of interest. Next, we developed a statistical method termed *GeneScore*, which is a weighted sum of the co-expression connectivity, and the weights are the *-log*_*10*_(P-values) from our differential expression analyses. As a normalization factor, we used the genomic control, which is the ratio of the observed median to the expected median test statistic^23^.

Among the 22 genes in the 16p11.2 locus that are expressed in cerebral organoids, 20 of these genes are also expressed in post-mortem brain samples from the BrainSpan Project. Similarly, when we calculated *GeneScores*_*all*_ using all genes detected from the 16p11.2 organoid RNA sequence data, we found that we were unable to prioritize any of the 11 genes in the 16p11.2 locus (P(*GeneScore*_*all*_)=0.71 to 0.97, **Fig. 3C, Supplementary Table 14**). Among the 13 genes in the 15q11-13 locus that are expressed in cerebral organoids, 11 of these genes are also expressed in post-mortem brain samples from the BrainSpan Project. We calculated *GeneScores*_*all*_ using all genes detected from the 15q11-13 organoid RNA sequence data, but were unable to prioritize any of the 11 genes in the 15q11-13 locus (P(*GeneScore*_*all*_)=0.38 to 0.39, **Fig. 3D, Supplementary Table 14**).

**Figure 2:**
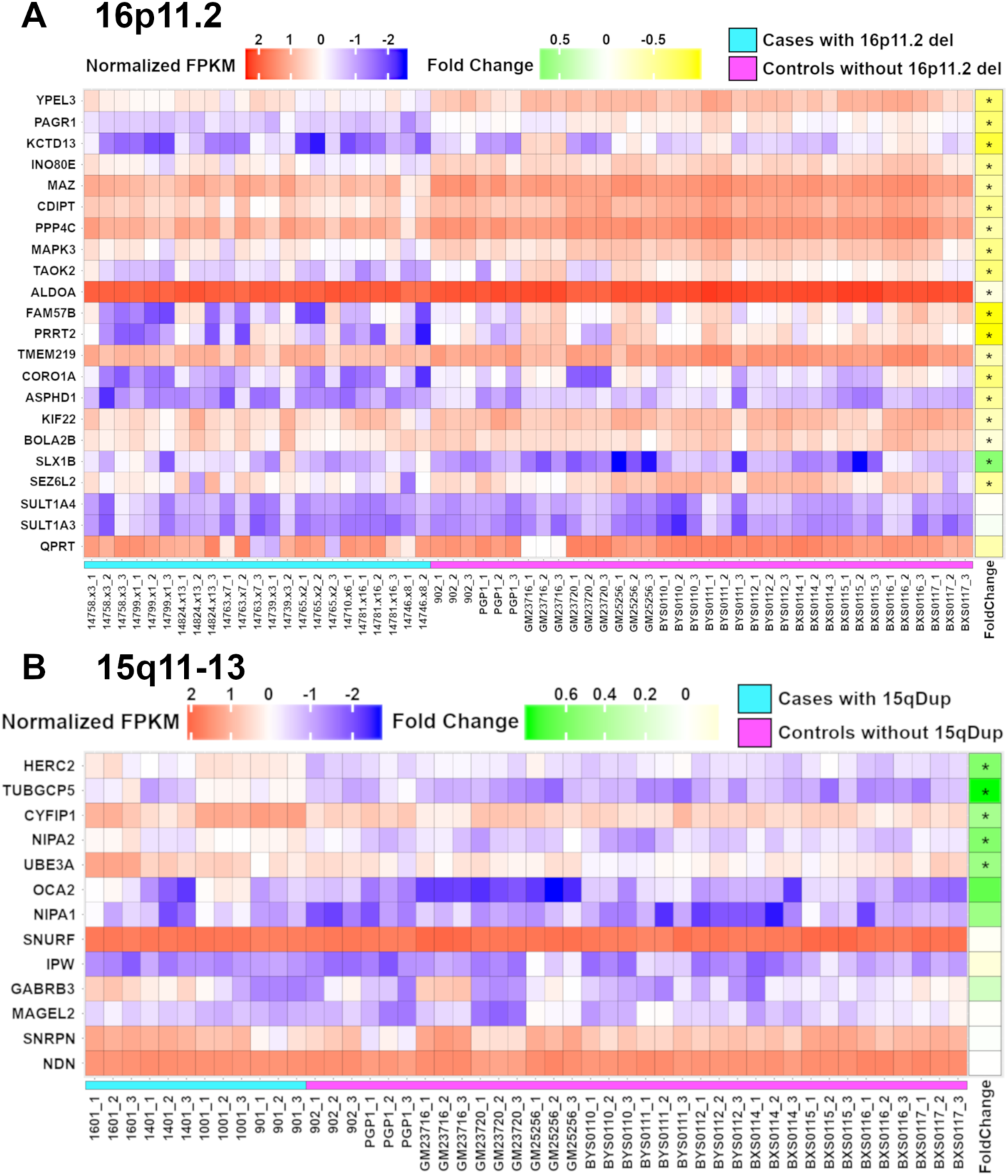
Expression of the genes in the 16p11.2 and 15q11-13 loci. **(A)** Heatmap representation of the normalized expression (FPKM) for all cases with 16p11.2 deletions (samples are indicated by the turquoise bar) and controls without the deletions (samples are indicated by the pink bar) across the 22 genes in the 16p11.2 locus. **(B)** Heatmap representation of the normalized expression (FPKM) for all cases with 15q11-13 duplications (samples are indicated by the turquoise bar) and controls without the duplications (samples are indicated by the pink bar) across the 13 genes in the 15q11-13 locus.

**Figure 3:**
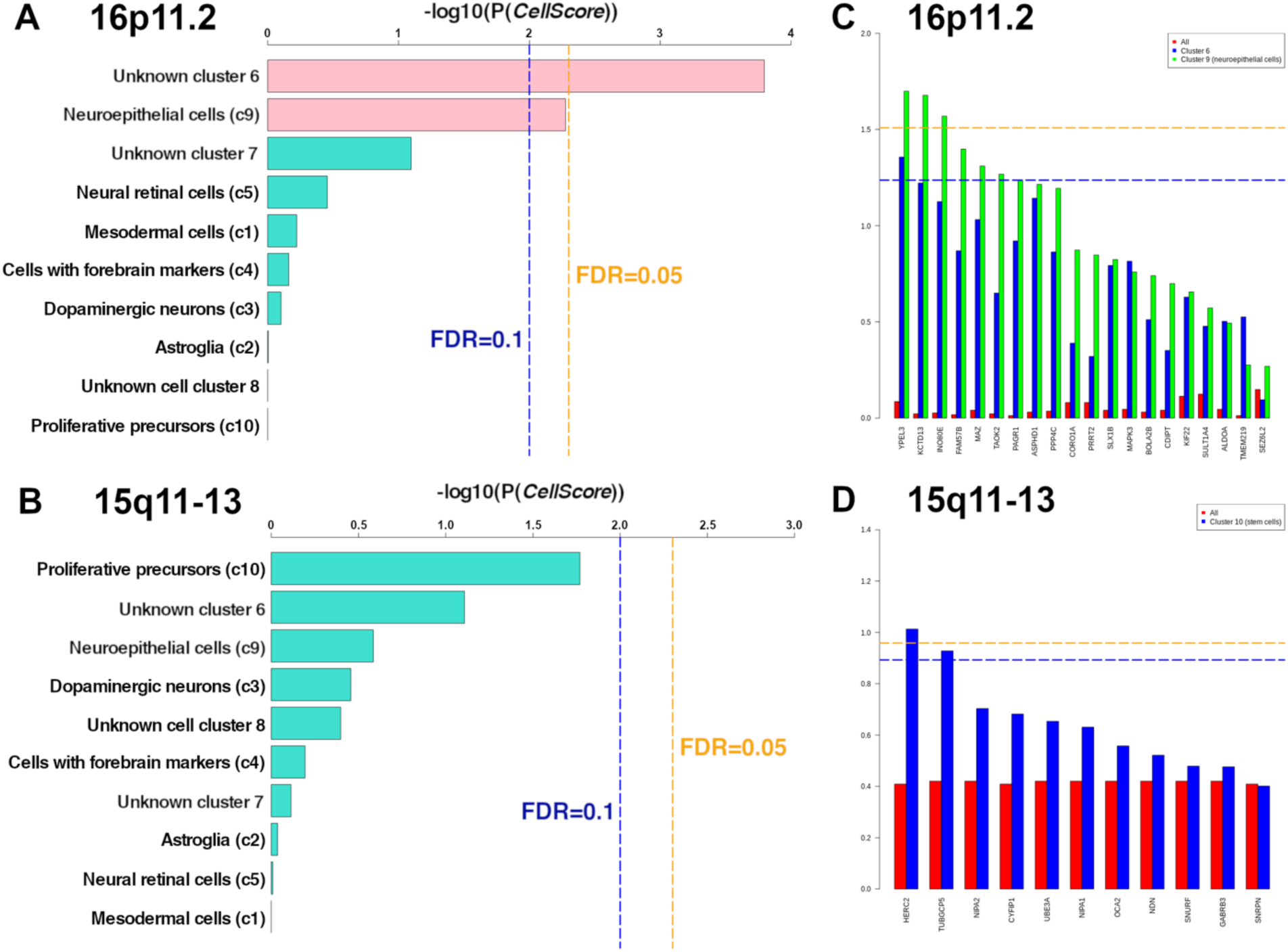
Prioritized critical cell types for the 15q11-13 and 16p11.2 locus. **(A)** Barplot showing the cell type results for the 16p11.2 locus, sorted in decreasing order of *- log*_*10*_(P(*CellScores*)), with the bars representing clusters that are likely to be critical cell types (FDR≤0.1) colored in pink and the rest of the clusters colored in turquoise. **(B)** Barplot showing the cell type results for the 15q11-13 locus, sorted in decreasing order of *-log*_*10*_(P(*CellScores*)), with the bars representing clusters that are likely to be critical cell types (FDR≤0.1) colored in pink and the rest of the clusters colored in turquoise. **(C)** Barplot showing the driver gene results for the 15q11-13 locus, sorted in decreasing order of *-log*_*10*_(P(*GeneScore*_*c9*_)), with red bars representing *-log*_*10*_(P(*GeneScore*_*all*_)), blue bars representing *-log*_*10*_(P(*GeneScore*_*c6*_)), and green bars representing *-log*_*10*_(P(*GeneScore*_*c9*_)). **(D)** Barplot showing the driver gene results for the 15q11-13 locus, sorted in decreasing order of *-log*_*10*_(P(*GeneScore*_*c10*_)), with red bars representing *-log*_*10*_(P(*GeneScore*_*all*_)), and blue bars representing *-log*_*10*_(P(*GeneScore*_*c10*_)).

### Cell type specific co-transcriptional network modeling can prioritize driver genes in 16p11.**2 and 15q11-13**

Given our earlier observation that cluster 9 comprising of neuroepithelial cells and cluster 6, are likely to important for the 16p11.2 locus, we hypothesized that we can obtain higher sensitivity to detect candidate driver genes by focusing on cell type specific signatures. When we adapted our *GeneScore* calculations to include only cell type specific genes that were identified in clusters 6 and 9, we found that 3 genes (*YPEL3* [MIM 609724], *KCTD13* and *INO80E* [MIM 610169]) were significantly prioritized as high-confidence candidate driver genes with FDR≤0.05 in cluster 9 (**Fig. 3C, Supplementary Tables 14-15**), and another 4 genes (*FAM57B* [MIM 615175], *MAZ* [MIM 600999], *TAOK2* and *PAGR1* [MIM 612033]) were prioritized as lower confidence candidate driver genes with FDR≤0.1 in cluster 9. Interestingly, we did not find any high-confidence candidate gene with FDR≤0.05 in cluster 6, and only 1 lower confidence candidate gene with FDR≤0.1 in cluster 6 (*YPEL3*).

One of the 3 high-confidence driver genes in cluster 9 (*KCTD13*) was initially implicated as an ASD risk gene that modulated brain sizes in zebrafish^7^, but recent studies using *KCTD13*-deficient mice and zebrafish did not observe any differences in brain sizes or neurogenesis^8,9^. Through the Orgo-Seq framework on patient-derived cerebral organoids, we found that *KCTD13* is one of 3 key driver genes in the 16p11.2 locus modulating the proportions of neuroepithelial cells in organoids with FDR≤0.05.

In *Ypel3*^-/-^ mice, an association with absence of startle reflex (P=2.2×10^−5^) and an association with short tibia (P=5.1×10^−6^) have been reported^24^. *TAOK2* and *MAZ* are targets of the fragile X mental retardation protein (FMRP)^25^; heterozygous and knockout mice for *TAOK2* have been recently reported to show impairments in cognition, anxiety and social interaction^18^, while mutations in *MAZ* decreases the promotor activity of NMDA receptor subunit type 1 during neuronal differentiation^26^.

Given that cluster 10 comprising of mainly stem cells was the top prioritized cell type for the 15q11-13 locus, we adapted our *GeneScore* calculations to include only cell type specific genes that were identified in cluster 10, and found *HERC2* and *TUBGCP5* as significant candidate driver genes at FDR≤0.1 (**Fig. 3D, Supplementary Tables 14-15**). Homozygous missense and deletion mutations in *HERC2* have been previously implicated in Amish individuals with severe neurodevelopmental disorders, with phenotypic features similar to Angelman Syndrome^22,27^. *HERC2* is also a key regulator of *UBE3A*^28^, which is another gene in the 15q11-13 locus, and mutations in *UBE3A* have been associated with ASD^19^. Rare mutations in *TUBGCP5* have also been reported in patients with ASD or intellectual disability^29^.

### CRISPR-edited mosaic organoids confirm enrichment of *KCTD13* mutants in neuroepithelial cells

To validate one of our results that *KCTD13* is a key driver gene in the 16p11.2 locus modulating the proportions of neuroepithelial cells in the patient-derived organoids, and to resolve prior conflicting results from *KCTD13*-deficient animal models^7-9^, we used a CRISPR-based approach^30-33^ to directly measure the effects of knockouts in cerebral organoids. We created *KCTD13* insertion and deletion mutations in iPSCs from a control individual (PGP1) using CRISPR/Cas9 with a synthetic guide RNA (gRNA). Next, we differentiated mosaic cerebral organoids from a mixture of iPSCs harboring *KCTD13* mutations (edited cells) and iPSCs with reference sequences (unedited cells). After 84 days, we harvested the mosaic cerebral organoids and dissociated single cells from the organoids for fluorescence activated cell sorting (FACS). We selected 4 antibody markers for FACS – NeuN for neuronal cells, Nestin for neural progenitor cells, TRA-1-60 for stem cells and mouse IgG2A as a negative control. DNA was extracted from the sorted cells and MiSeq sequencing was performed to identify the proportions of *KCTD13* mutations in these sorted cell populations.

This validation approach allows us to test our observed results from RNA sequence data obtained from the patient-derived cerebral organoids using an orthogonal approach with cell type specific protein markers on the CRISPR-edited mosaic cerebral organoids. If *KCTD13* mutations do not affect a specific cell population, we expect to observe that the mutations are not significantly enriched in cells that are positive or negative for that cell type marker, as illustrated in **Fig. 4A**. However, if *KCTD13* mutations affect a specific cell population, we expect to observe that the mutations are significantly enriched in the cells that are positive for that cell type marker.

**Figure 4:**
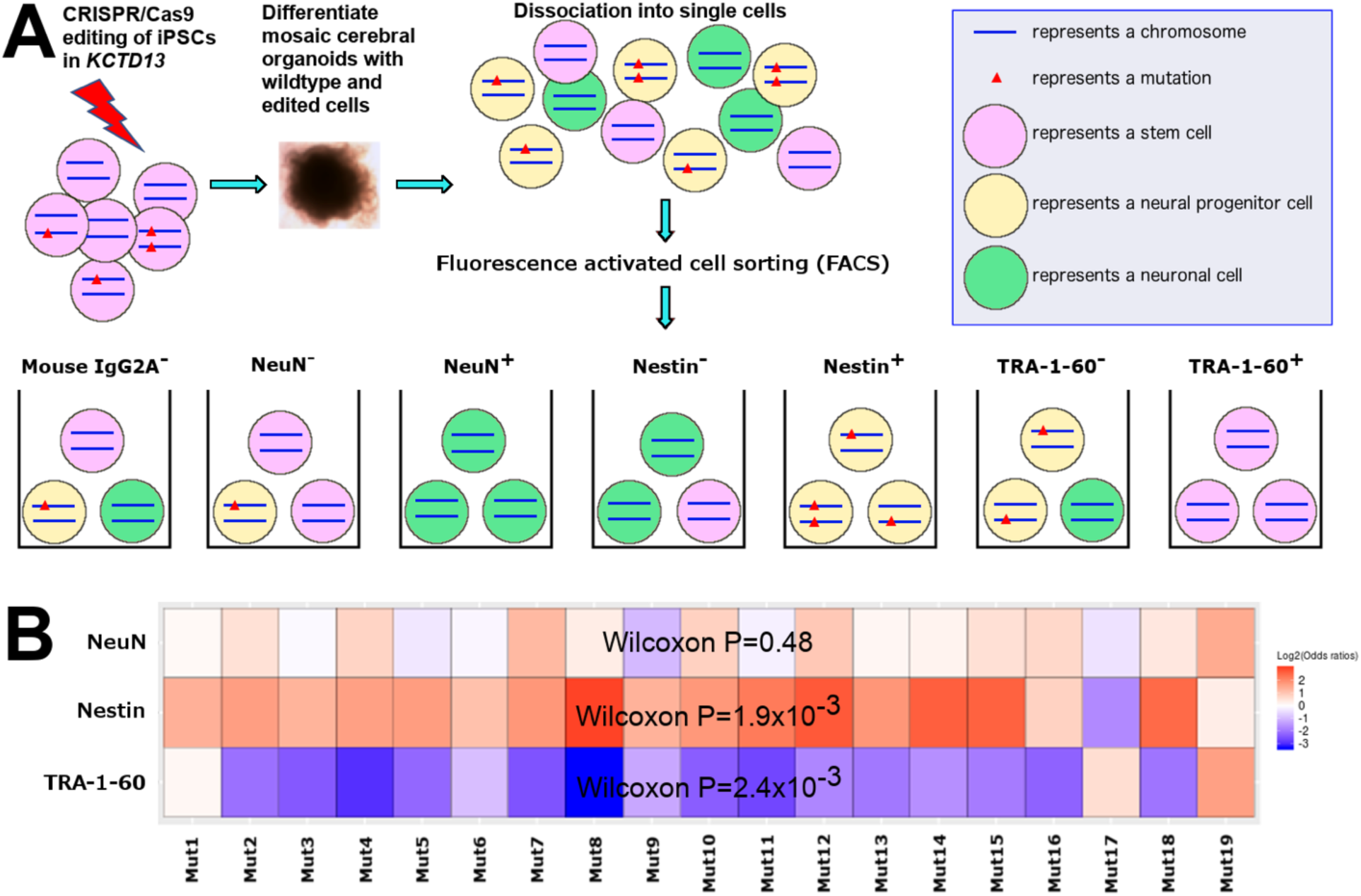
FACS-based framework to identify cell types affected by *KCTD13* mutations in CRISPR-edited cerebral organoids. **(A)** Schematic of our validation framework where we performed CRISPR/Cas9 editing in the *KCTD13* locus on iPSCs (denoted as pink circles) from a control individual and pooled a mixture of edited and unedited cells to differentiate cerebral organoids. These mosaic organoids, comprising of different cell types (represented as differently colored circles) with different genotypes in *KCTD13*, were subsequently dissociated into single cells for FACS into 7 sorted pools of cells. **(B)** Heatmap representations of the proportions of the KCTD13 mutations in NeuN^+^ cells compared to NeuN^-^ cells (first row), Nestin^+^ cells compared to Nestin^-^ cells (second row) and TRA-1-60^+^ cells compared to TRA-1-60^-^ cells (third row). Red represents an enrichment of these mutations in sorted cells that are positive for the respective cell type marker, while blue represents an enrichment of these mutations in sorted cells that are negative for the respective cell type marker.

Our results from the CRISPR-edited mosaic organoids suggest that *KCTD13* mutations resulted in increased numbers of neuroepithelial cells marked by Nestin (Wilcoxon P=4.6×10^−3^), and are depleted in stem cells marked by TRA-1-60 (Wilcoxon P=3.5×10^−3^), as shown in **Fig. 4B** and **Supplementary Table 16**. There is no statistically significant enrichment of *KCTD13* mutations in neurons marked by NeuN (Wilcoxon P=0.47), suggesting that *KCTD13* mutations primarily affect neural progenitor cells but not neurons. Using a more stringent subset of mutations that map uniquely to the *KCTD13* locus, we similarly observed that there is a significant increase in neuroepithelial cells with *KCTD13* mutations (Wilcoxon P=1.9×10^−3^), decrease in stem cells with *KCTD13* mutations (Wilcoxon P=2.4×10^−3^), but no difference in the number of neurons with *KCTD13* mutations (Wilcoxon P=0.48).

These results confirm our earlier findings that mutations in *KCTD13* in the 16p11.2 locus lead to increased proportions of neural progenitor cells in human cerebral organoids, and these results also agree with clinical observations that patients with 16p11.2 deletions have increased brain sizes, marked by increased proportions of neural progenitor cells^5,6^. Orgo-Seq provides a quantitative framework for identifying cell type specific driver genes and the results can be rapidly followed up using dissociated single cells from CRISPR-edited mosaic organoids.

## DISCUSSION

Cerebral organoids are an emerging human-derived model system for studying complex neurodevelopmental and neuropsychiatric disorders^2,34-38^. These cerebral organoids comprise of many different cell types, so this effectively allows us to test multiple hypotheses in multiple cell types that were differentiated under the same conditions. Given the differences between the human brain and brains from model organisms, as well as the challenges in obtaining large numbers of human post-mortem brain tissue with specific disease-associated mutations, it will be increasingly important to develop technologies and methods to utilize patient-derived human cerebral organoids as a model system for studying complex neurodevelopmental and neuropsychiatric disorders^3^, as well as human-specific brain evolution^39^. In addition, a major strength of using patient-derived organoids instead of CRISPR-edited organoids for discoveries is that the patient-derived organoids can model the diverse genetic backgrounds found in humans, and are more representative of tissue derived from different patients. As such, technology and methods development to enable unbiased high-throughput discoveries using patient-derived organoids can leverage on the complexity of human genetics for making important discoveries in disease biology.

In our work, we describe the Orgo-Seq framework to allow the identification of cell type specific driver genes from patient-derived cerebral organoids that are important in ASD-associated CNVs, by utilizing both bulk transcriptomics and single-cell transcriptomics data. Orgo-Seq allows us to overcome technical limitations such as capture efficiencies with detecting critical cell types and cell type specific driver genes using single-cell RNA sequencing alone. Our framework can be generalized for identifying specific types of neurons or other cell types of interest, as well as cell type specific driver genes for many other CNVs that have been robustly associated with complex neurodevelopmental and neuropsychiatric disorders^4,40^. It is interesting to note that we were unable to prioritize any candidate driver genes when using co-expression patterns of all genes whose expression were detected in the cerebral organoids, but we were able to identify cell type specific candidate driver genes when using co-expression patterns of cell type specific genes. Studying cell type specific driver genes and processes will be increasingly important for complex diseases^41^, and cerebral organoids can be powerful model systems that allow us to identify cell type specific processes in complex diseases using a genotype-driven approach.

The Orgo-Seq framework is scalable and generalizable, and enables researchers who are interested in Mendelian or complex diseases to discover cell types and cell type specific driver genes from organoids, not limited to the brain, by utilizing bulk RNA sequencing and leveraging on ongoing large-scale single-cell RNA sequencing studies that are being performed on multiple tissue and cell types from multiple species^41,42^, as well methods development for the integration of multiple single-cell RNA sequence datasets^39,43^. In addition, the Orgo-Seq framework allows us to reanalyze the bulk RNA sequence data that we have already generated from the patient-derived organoids for making new discoveries about cell types and cell type specific driver genes, by using new large-scale single-cell RNA sequencing data that have been, or will be, generated on large numbers of single cells^42^, or single-cell RNA sequencing data that will be generated using new spatial-informative technologies^44,45^.

We have also demonstrated a validation approach to rapidly create mosaic cerebral organoids from a mixture of edited and unedited cells, and identify cell types affected by these mutations using cell type specific antibodies. A major strength of mosaic cerebral organoids differentiated from a mixture of edited and unedited iPSCs is that similar conditions are maintained for all the edited and unedited cells across different cell types, given that all cells are differentiated within and dissociated from the same organoids. As such, we can leverage on the heterogeneity of the cerebral organoids for creating a self-controlled mixture of cells for validating hypotheses about cell types affected by disease-associated mutations.

In summary, we have established a set of quantitative frameworks for generating and validating hypotheses about cell type specific driver genes involved in complex neurodevelopmental and neuropsychiatric disorders using a human-derived model system.

## METHODS

### CNV analyses

A total of 25 iPSCs (1 clone per iPSC) were obtained as from Coriell Institute, ATCC, Harvard Stem Cell Institute and Simons VIP collection. All iPSCs and cerebral organoids were tested negative for mycoplasma using the LookOut Mycoplasma PCR Detection kit (Sigma MP0035). iPSCs from all donors were passaged until they were confluent, and 2 million cells per donor were counted using an automated cell counter, and washed twice in 1x DPBS, before flash freezing the cell pellets. The frozen cell pellets were sent on dry ice to Cell Line Genetics, where genomic DNA was extracted from the cells, and quality control was performed using Nanodrop, Qubit and agarose gel analyses. The Agilent 60k standard aCGH was used to identify CNVs, and the CNVs were compared to the Database of Genomic Variants (CNV-DGV_hg19_May2016) to identify CNVs that are common in the general population (**Supplementary Table 2**). All 4 donors with 15q11-13 duplications were confirmed to harbor the duplications, all 9 donors with 16p11.2 deletions were confirmed to harbor the deletions, and all 12 control individuals were confirmed not to harbor any duplication in the 15q11-13 locus, or deletion in the 16p11.2 locus.

To identify smaller exonic CNVs, we further performed CNV analyses from whole-exome sequence data on all donor iPSCs. DNA was extracted from iPSC cell pellets for all donors using the standard protocol for AccuPrep Genomic DNA Extraction Kit (Bioneer K-3032), and Nanodrop was used to evaluate the quantity and quality of the extracted DNA samples. 1μg of DNA per iPSC was sent on dry ice to Macrogen, where quality control was performed using Quant-iT PicoGreen dsDNA Assay Kit (Life Technologies P7589) with Victor X2 fluorometry, and the Genomic DNA ScreenTape assay (**Supplementary Table 1**). The DNA Integrity Number (DIN) threshold used for exome sequencing was 6, and the mean DIN across all control samples was 8, the mean DIN across all samples with 15q11-13 duplications was 7.7 and the mean DIN across all samples with 16p11.2 deletions was 7.9, but there were no significant differences between the DNA quality from the iPSCs with 15q11-13 duplications versus the control iPSCs (two-sided Wilcoxon P=0.2), or the iPSCs with 16p11.2 deletions versus the control iPSCs (two-sided Wilcoxon P=0.62). The Agilient SureSelect V5-post kit was used for capture and the library was sequenced using NovaSeq 6000 (150 paired end). CNV calling on the exome sequence data was performed using CoNIFER^46^, and all exonic CNVs detected from the iPSCs are shown in **Supplementary Table 3**. Among the cases with 15q11-13 duplications or 16p11.2 deletions, only a smaller deletion in the 16p11.2 locus encompassing exons in *TAOK2* and an intron in *BOLA2B* was found detected from the whole exome sequence data for proband 14824.x13.

### Cerebral organoid differentiation

We adapted our cerebral organoid differentiation protocol according to a previously described protocol^2^ (**Supplementary Fig. 1A**). For embryoid body formation, cells were counted using an automated cell counter and 900,000 iPSCs were re-suspended in 15ml of mTeSR medium (Stemcell Technologies 85850) with 50μM ROCK inhibitor (Santa Cruz sc-216067A), and 150μl was seeded into individual wells of a 96-well ultra-low attachment Corning plate (ThermoFisher CLS7007). On Day 6, 50μl of mTeSR medium with a single embryoid body was transferred to individual wells of 24-well ultra-low attachment Corning plates (ThermoFisher CLS3473) with 500μl of neural induction media per well. On Day 8, another 500μl of neural induction media was added to each well of the 24-well plates. On Day 10, a droplet comprising of 10μl of neural induction media with an organoid was placed onto a single dimple on Parafilm substrate, and 40μl of Matrigel (Corning 354234) was added to each organoid to encapsulate it. The Matrigel droplets were incubated at 37°C for 15 minutes before they were scrapped into single wells of the 24-well plates using a cell scraper. 1ml of differentiation media with 10% penicillin streptomycin (ThermoFisher 15140122) per well was used to passage the organoids every 2-4 days, and the plates of organoids were placed on an orbital shaker at 90rpm in the incubator. A previous publication noted that bioreactor-related growth environment is a key factor in controlling cell type identity from organoids to organoids^1^, and similarly, we had observed batch effects in the rates of cell death while differentiating multiple organoids in the same well of multi-well plates. As such, we differentiated single organoids in individual wells of the 24-well plates, to minimize batch effects for individual organoids due to the growth environment.

### Cerebral organoid cryosection and immunostaining

Cerebral organoids were rinsed twice with 1× DPBS, fixed in 4% paraformaldehyde at 4°C for 30-60 minutes, immersed in 30% sucrose overnight, embedded in optimal cutting temperature compound (OCT), and 8-micron sections are collected with a cryostat. Cryosections of fixed cerebral organoids were immunostained with antibodies against Sox2 (Santa Cruz sc-17320), Tbr2 (Abcam ab-23345), Tuj1 (Covance MMS-435P) and Alexa Fluor secondary antibodies (ThermoFisher).

### RNA extraction, sequencing, alignment and annotation

It was previously noted that some cell types are found in only 32-53% of organoids, using single-cell RNA sequencing^1^. In order to reduce variability across replicates, as well as to obtain sufficient representation of all cell types, we pooled 20 separate organoids from different wells and different plates, as one replicate. The organoids in each replicate were pelleted at 1,000*g* for 1 minute, and the supernatant was removed, before washing twice in DPBS. RNA from 1-3 replicates was extracted for each individual (**Supplementary Table 1**). The organoids were homogenized using mechanical disruption in lysis buffer, and RNA extraction was performed using the PureLink RNA Mini Kit (ThermoFisher 12183018A), according to the manufacturer’s protocol. RNA samples were treated with Ambion DNase I (ThermoFisher AM2222) according to the manufacturer’s protocol, before they were frozen and sent on dry ice to Macrogen.

At Macrogen, DNA quantity was measured using Quant-iT PicoGreen dsDNA Assay Kit (Life Technologies P7589) with Victor X2 fluorometry, and RNA quantity was measured using Quant-iT RiboGreen RNA Assay Kit (Life Technologies R11490). The RNA Integrity Number (RIN) was measured using an Agilent Technologies 2100 Bioanalyzer or TapeStation, and the RIN value threshold used was 6 (**Supplementary Table 1**). Ribosomal RNA depletion using TruSeq Stranded RNA with Ribo-Zero (Human) and paired-end 101bp sequencing with at least 30 million reads per sample was performed. Library size checks were performed using an Agilent Technologies 2100 Bioanalyzer or TapeStation, and quantification of the libraries was performed according to the Illumina qPCR quantification guide. Reads were trimmed using Trimmomatic 0.32, then mapped to the hg19 human genome sequence using TopHat 2.0.13, and transcript assembly was performed using Cufflinks 2.2.1 to calculate the fragments per kilobase per million reads (FPKM) values for each transcript. In addition, the reads were mapped to the hg19 sequence using STAR 2.4.0f1, and single nucleotide variant calling on the aligned sequences was performed using GATK 3.3-0 HaplotypeCaller. Annotation for the single nucleotide variants was performed using SeattleSeq Annotation 138, and single nucleotide variants detected from the RNA sequence data were compared between replicates from the same individual and verified for concordance (*r*>0.95), to ensure that there was no sample mix-up.

### Data processing and quality control

The mean RIN values for the control samples, 15q11-13 samples and 16p11.2 samples were 7.9, 8.1 and 8.2 respectively (**Supplementary Table 1**). We performed a two-sided Wilcoxon rank sum test between the RIN values for the control samples versus the 15q11-13 samples, but did not observe significant differences (P=0.43). Similarly, we did not observe significant differences between the RIN values for the control samples versus the 16p11.2 samples (P=0.13). Neither did we observe significant differences between the RIN values for the 15q11-13 samples versus the 16p11.2 samples (P=0.47).

After selecting the transcript with the highest mean FPKM across all samples (including all cases and controls) for each gene, there were 25,727 unique transcripts or genes. We further performed quality control to remove genes that were not expressed, or had high intra-individual or inter-individual variance. Genes that were not expressed in the cerebral organoids (mean FPKMs across all samples < 2) were removed, resulting in a smaller set of 11,300 genes. We calculated the mean FPKMs across all samples, including all case and control samples. However, we used only the control samples for calculating the standard deviations in gene expression, to preserve genes that truly contribute to biological variation between the case and control organoids. Inverse rank sum normalization was performed on the expression values that were subsequently used in the downstream analyses, as the normalization procedure reduces outlier expression values.

There were 860 genes with more than 2 standard deviations in any intra-individual variance calculated across the control samples (**Supplementary Fig. 2A, Supplementary Table 4**), and 869 genes with more than 1.5 standard deviations in inter-individual variance calculated between the control samples (**Supplementary Fig. 2B, Supplementary Table 4**), resulting in a total of 1,322 unique outlier genes. After removing all outlier genes with high variability, there are a total of 9,978 unique genes. Pairwise Pearson’s correlations (*r*^2^) were performed for each pair of replicates from an individual to calculate the intra-individual correlations, and each pair of replicates from different individuals to calculate the inter-individual correlations.

### Comparing BrainSpan samples with cerebral organoid samples

The BrainSpan project (http://www.brainspan.org) provides a high-resolution map of 22,326 genes detected using RNA sequencing on 578 post-mortem brain samples from various brain regions in prenatal brains (8 pcw) to adult brains (40 years old)^12^. We downloaded the “RNA-Seq Gencode v10 summarized to genes” dataset from the BrainSpan Project for our analyses (http://www.brainspan.org/static/download.html). For comparing RNA sequence data from prenatal brain samples from the BrainSpan Project with RNA sequence data from cerebral organoids, we included only brain regions where more than 50% of samples were available for those regions (≥9 samples). We performed two-sided Wilcoxon rank-sum test to evaluate if the mean Pearson’s correlations between the organoids and prenatal brain samples were significantly higher than the mean Pearson’s correlations between the organoids and postnatal brain samples. We further calculated Pearson’s correlations for each pair of genes from the BrainSpan RNA sequence data.

We observed that after removing highly variable genes, the Pearson’s correlations between RNA sequence data from the organoids with the 578 post-mortem brain samples ranged from 0.21 to 0.82 (**Supplementary Fig. 5A**). Prior to removing highly variable genes, the Pearson’s correlations between RNA sequence data from the organoids with the post-mortem brain samples ranged from 0.14 to 0.93 (**Supplementary Fig. 5D**). The larger variance in correlations prior to removing highly variable genes was primarily driven by high outlier correlations in all replicates from two control samples (BYS0110 and BXS0115). For instance, the mean Pearson’s correlation between all organoid samples excluding those differentiated from BYS0110 and BXS0115, with cerebellar cortex from a 16pcw fetal brain sample is 0.33. However, the mean Pearson’s correlations between the 16pcw fetal brain sample with organoid samples from BYS0110 is 0.84 and with organoid samples from BXS0115 is 0.92.

### Differential gene expression analyses

Principal components analyses on all samples across control individuals without deletions or duplications, and individuals with 16p11.2 deletions or 15q11-13 duplications showed that PC1 alone accounted for 88% of the variance in gene expression (**Supplementary Fig. 3A**). We performed differential expression analyses using linear regression in R (`lm` function), with PC1 as a covariate, and performed multiple hypotheses correction using the Benjamini-Hochberg false discovery rate in R (`p.adjust`), shown in **Supplementary Tables 5-7** and **Supplementary Table 9**. Given the relatively small number of samples used in our study^47^, and since PC1 captures 88% of the variance in gene expression and is a surrogate factor for several sample variables, we included only PC1 as a covariate in our linear regression analyses to identify differentially expressed genes. We further plotted the first 2 principal components between control individuals without deletions or duplications, and individuals with 16p11.2 deletions or 15q11-13 duplications, but did not observe major stratification between the cases and controls in the first 2 principal components (**Supplementary Fig. 6**). Given that the inter-individual correlations observed between samples from different individuals are similarly high compared to the intra-individual correlations observed between replicates from the same individual, and given the relatively small number of individuals in our study that limits the number of permutations, we performed linear regression using all samples as independent samples. We had also performed bulk RNA sequencing on 1-3 replicates for each individual, to ensure that the results were not skewed by RNA sequence data from a few outlier individuals.

### Permutation schemes for 16p11.**2 SetA and 15q11-13**

We permuted the case-control status of each organoid replicate to obtain null distributions. However, given the relatively small numbers of samples, we wanted to avoid creating permuted instances where the permuted cases are actual case samples and the permuted controls are actual control samples. Differential expression analyses on these permuted instances will result in the detection of true biological differences, instead of creating a baseline non-biological measurement for the null distribution. As such, we developed a permutation strategy by sampling permuted case samples from the actual control samples only (**Supplementary Fig. 7**). Furthermore, to ensure that we have the same numbers of case and control samples in our permutations, as the numbers of case and control samples from our actual experiments, we assigned all the actual case samples to be permuted control samples. We refer to the cases in the permutations as “pseudo-cases”, and the controls in the permutations as “pseudo-controls”.

For 16p11.2 SetA, we performed differential expression analyses for 23 samples differentiated from all individuals with 16p11.2 deletions (cases) versus 36 samples differentiated from unaffected controls without the deletion (controls). To obtain a null distribution, we randomly assigned 23 samples from the 36 control samples as pseudo-cases, and assigned the initial 23 samples, together with the remaining control samples as pseudo-controls, for 100,000 permutations. Subsequently, we performed linear regressions with PC1 as a covariate on all the expression data for the 100,000 permutations.

For the 15q11-13 results, we performed differential expression analyses for 12 samples differentiated from individuals with ASD and 15q11-13 duplications (cases) versus 36 samples differentiated from unaffected controls without the duplications (controls). To obtain a null distribution for comparing the observed statistics, we randomly assigned 12 samples from the 36 control samples as pseudo-cases, and assigned the initial 12 samples, together with the remaining control samples, as pseudo-controls, for 100,000 permutations. Subsequently, we performed linear regressions with PC1 as a covariate on all the expression data for the 100,000 permutations.

### Calculation of *CellScore* and P(*CellScore*)

There are 10 major clusters of cell types identified using unbiased clustering on single-cell RNA sequence data from organoids, and each cell cluster has an associated list of genes identified using Drop-seq, and was assigned a cell cluster identity using previously published data from homogeneous cell populations^1^. We downloaded the data from Quadrato *et al*., and observed that in these full lists of cluster genes, there are some genes that are present in multiple cell clusters, and that these genes are not cell type specific. To enrich for cell type specific genes, we further identified a smaller subset of genes that are uniquely found in each cell type cluster but are not present in other cell type clusters, which we termed as “cell type specific genes” (**Supplementary Table 10**). We termed the genes that are found in multiple cell clusters as “non-cell type specific genes”.

We calculated *CellScore* for each cluster by summing up the -*log*_*10*_-transformed P-values from the differential expression results for each gene *y*in the cluster (*P*_*y*_), divided by the total number of genes in the cluster (*Num*_*y*_), and obtained the difference between the calculated *CellScores* for the specific genes versus the non-specific genes, where *P*_*specific_y*_ is the P-value of each cell type specific gene in the cluster, *Num*_*specific_y*_ is the number of cell type specific genes in the cluster, *P*_*non-specific_y*_ is the P-value of each non-cell type specific gene in the cluster, and *Num*_*non*-*specific_y*_ is the number of non-cell type specific genes in the cluster. Taking the difference between the calculated *CellScores* for the cell type specific genes versus the non-cell type specific genes allows us to obtain a normalized *CellScore* that is adjusted for other inherent factors that can similarly affect the expression of non-cell type specific genes.

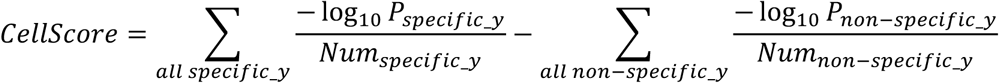

We obtained a null distribution for *CellScore* by performing 100,000 permutations (see Permutation schemes for 15q11-13 and 16p11.2 SetA), and performed linear regressions for each permutation. Next, we estimated the probability of the observed *CellScore* for each cluster by comparing with the null distribution (*CellScore*_*permuted*_):

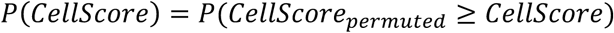

To identify significant clusters, we calculated an FDR threshold of 0.05 after Bonferroni correction for multiple hypotheses, i.e. P = 0.05/10 = 0.005; and similarly, for an FDR threshold of 0.1, or P = 0.1/10 = 0.01.

### Analysis of post-mortem brain samples with 15q11-13 duplications

The differential expression results on post-mortem brain samples from the cortex with or without 15q11-13 duplications were downloaded from a prior publication^10^. We calculated *CellScores* for each of the 10 cell type clusters by using the P-values from the differential expression results for each gene. To calculate P(*CellScore*), we compared the observed *CellScore* from the post-mortem brain samples against the null *CellScore* distributions for each of the 10 cell type clusters generated by the permutations using the expression data from the cerebral organoids with 15q11-13 duplications, accounting for the precise numbers of genes used in the calculations of *CellScores* from the post-mortem brain samples. We calculated a weighted average P-value for the results from the cerebral organoids with 15q11-13 duplications and the results from the post-mortem brain samples with 15q11-13 duplications (**Supplementary Table 12**), which allows us to evaluate the combined *CellScore* results from the cerebral organoids and the post-mortem brain samples.

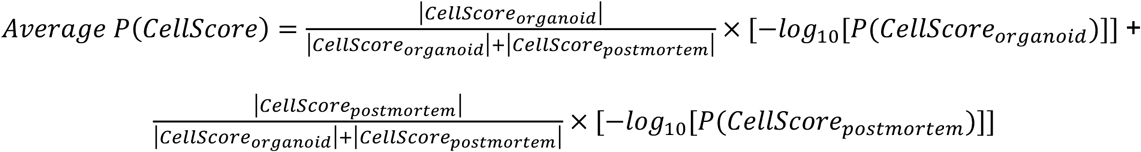

where |*CellScore*_*orangnoid*_ and |*CellScore*_*postmortem*_| are the absolute *CellScore* values calculated from the cerebral organoids and post-mortem brain samples respectively, and *P*(*CellScore*_*orangnoid*_) and *P*(*CellScore*_*postmortem*_) are the P(*CellScore*) values calculated from the cerebral organoids and post-mortem brain samples respectively.

### Calculation of *GeneScore* and P(*GeneScore*)

There were 22 genes in the 16p11.2 locus that are expressed in the cerebral organoids, but 2 of the genes (*SULT1A3* and *QPRT*) were not found in the BrainSpan expression dataset, and were excluded from our candidate driver gene analyses. Similarly, there were 13 genes in the 15q11-13 locus that are expressed in the cerebral organoids. However, 2 of the genes (*IPW* and *MAGEL2*) are not found in the BrainSpan expression dataset, and were excluded from our candidate driver gene analyses using *GeneScore*.

We calculated *GeneScore* for each gene *x* in a CNV locus using the total sum of the Pearson’s correlation 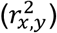 of gene *x* with each gene *y*in the BrainSpan Project^48^, multiplied by the -*log*_*10*_-transformed P-values from the organoid differential expression results for gene *y* (*P*_*y*_), and divided the scores by the total number of genes (*Num*_*y*_) from the BrainSpan Project with correlations available for gene *x*.

We obtained a null distribution for *GeneScore* by performing 100,000 permutations (**Supplementary Fig. 7**), and performed linear regressions on the expression data for each permutation. Next, we calculated *GeneScore* for each gene *x* based on the permuted linear regression results. Since our observation and each permutation comprises of different combinations of individuals who have been assigned as pseudo-cases or pseudo-controls, we calculated a representative statistic (genomic control or *λ*)^23^, which is the ratio of the observed median to the expected median test statistic, to evaluate the P-value distribution in each permutation, and normalized the observed and permuted *GeneScores* with the inverse of *log*_10._*λ*:

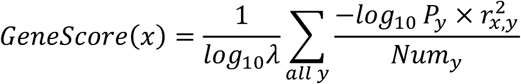

We estimated the probability of the observed *GeneScore* for each gene *x* by comparing the observed *GeneScore* with the null distribution (*GeneScore*_*permuted*_):

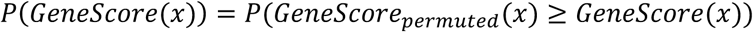

To evaluate the cell type specific *GeneScores*, we used the differential expression results from the same 100,000 permutations and calculated cell type specific *GeneScore* using only the specific and non-specific genes in each cell cluster (c1-c10). To estimate the FDR for the cell type specific *GeneScores* in the 16p11.2 and 15q11-13 loci, we sorted all the P-values calculated for the *GeneScores* from all clusters for each locus, to obtain the distributions of P-values. For each locus, we used the 5^th^ percentile P-value as the FDR threshold of 0.05, and 10^th^ percentile P-value as the FDR threshold of 0.1.

### CRISPR/Cas9-editing of cerebral organoids

The iPSCs used for CRISPR/Cas9-editing were from an unaffected control individual (PGP1). iPSCs were passaged until they were 50-75% confluent prior to nucleofection. Nucleofection was performed using the HSC-1 kit and B-016 protocol on an Amaxa nucleofector. Four gRNAs for *KCTD13* and Cas9 protein were ordered from Synthego, and we evaluated the efficiencies for on-target editing of *KCTD13* for each of the 4 gRNAs, as well as a combination of all 4 gRNAs, in iPSCs using nucleofection followed by MiSeq sequencing. We selected the gRNA with the highest on-target editing efficiency and the sequence is shown below.

*KCTD13* gRNA (chr16:29923312-29923331): UGAGGAUUGUACCAAAGUGA After nucleofection, the iPSCs were passaged for 7 days until they were confluent, and DNA extraction was performed on half of the iPSCs, followed by PCR and MiSeq sequencing to confirm the presence of locus-specific insertions and deletions (protocols and primers described below). 900,000 cells from the other half of the iPSCs were then differentiated into mosaic cerebral organoids.

### Preparation of mosaic cerebral organoids for antibody staining and FACS

Mosaic cerebral organoids were harvested after 84 days, and washed twice using 1× DPBS. 0.25% Trypsin-EDTA (ThermoFisher 25200056) was added to dissociate the cells for 30 minutes at 37°C on a shaking heat block at 300 rpm, before inactivating the trypsin using mTeSR medium and washed twice using 1× DPBS. To remove residual Matrigel, dissociated cells were filtered through 30 µm cell strainers (Miltenyi Biotec 130-041-407). Cells were counted using an automated cell counter and 12 million cells were fixed and permeabilized using equal volumes of 1% paraformaldehyde and permeabilization buffer (DPBS, 0.02% sodium azide, 2% FBS and 0.1% saponin) for 45 minutes. 3 million cells were used for each antibody and staining was performed for an hour, followed by FACS. The antibodies used were commercially available and Alexa Fluor 488-conjugated: mouse IgG2A control (R&D Systems IC003G), NeuN (Novus Biologicals NBP-92693AF488), Nestin (R&D Systems IC1259G) and TRA-1-60 (Novus Biologicals NB100-730F488). Cells that were negative for the mouse IgG2A antibody were collected as positive controls, and cells that were negative or positive for NeuN, Nestin or TRA-1-60 were collected.

### DNA extraction, PCR and MiSeq sequencing

Cells were washed twice with 1× DPBS and DNA extraction was performed using the standard protocol for AccuPrep Genomic DNA Extraction Kit (Bioneer K-3032). Locus-specific PCR was performed using the standard protocol for Q5 Hot Start Master Mix (New England BioLabs M094S). PCR was also performed on unedited DNA extracted from PGP1 iPSCs as a control for background.

The following primers were used for sequencing:

Forward primer: 5’-TTCTCTCGTACTCTCCGGCA -3’

Reverse primer: 5’-GCCTGGGCAACATAGTGAGA -3’

Barcoding was performed using Nextera indexes, followed by DNA clean-up using the Monarch PCR & DNA Cleanup Kit (New England BioLabs T1030S). Library preparations were quantified using the using KAPA library quantification kit (Kapa Biosystems KK4824) and pooled in equal concentrations prior to MiSeq v3 sequencing with 15% phiX control spike-in (Illumina FC-110-3001).

### Analyses on MiSeq data from CRISPR-edited organoids

Unique sequences detected from the MiSeq sequencing data were counted for each of the 8 samples (Mouse IgG2A^-^, NeuN^-^, NeuN^+^, Nestin^-^, Nestin^+^, TRA-1-60^-^, TRA-1-60^+^ and PGP1 unedited cells). To identify mutations that were specific to the CRISPR-edited cells but were not present in PGP1 unedited cells, sequences with less than 25 reads in PGP1 cells and had at least 25 reads in each of the 7 edited samples, were identified for further analyses (**Supplementary Table 16**). The number of reads for each mutant sequence was divided by the number of reads for the reference sequence in each sample to obtain normalized ratios. Two-sided Wilcoxon ranked sum test was performed to test the normalized ratios between the NeuN^-^ and NeuN^+^, Nestin^-^ and Nestin^+^, TRA-1-60^-^ and TRA-1-60^+^ samples. Since we are performing 3 Wilcoxon ranked sum tests across the 3 sets of cell types, the Bonferroni-corrected P-value threshold used was 0.05/3 = 0.017. Odds ratios were further calculated for each mutant sequence by dividing the normalized ratios for NeuN^+^, Nestin^+^ or TRA-1-60^+^ by the normalized ratios for NeuN^-^, Nestin^-^ or TRA-1-60^-^ samples respectively. Heatmaps to visualize the odds ratios for NeuN, Nestin and TRA-1-60 across all mutant sequences were plotted using `ggplot2` in R.

